# Metabolic differentiation of co-occurring Accumulibacter clades revealed through genome-resolved metatranscriptomics

**DOI:** 10.1101/2020.11.23.394700

**Authors:** Elizabeth A. McDaniel, Francisco Moya-Flores, Natalie Keene Beach, Pamela Y. Camejo, Ben O. Oyserman, Matthew Kizaric, Eng Hoe Khor, Daniel R. Noguera, Katherine D. McMahon

**Author notes:** equal contribution (Elizabeth A. McDaniel, Francisco Moya-Flores).

## Abstract

Natural microbial communities consist of closely related taxa that may exhibit phenotypic differences and inhabit distinct niches. However, connecting genetic diversity to ecological properties remains a challenge in microbial ecology due to the lack of pure cultures across the microbial tree of life. *‘Candidatus* Accumulibacter phosphatis’ is a polyphosphate-accumulating organism that contributes to the Enhanced Biological Phosphorus Removal (EBPR) biotechnological process for removing excess phosphorus from wastewater and preventing eutrophication from downstream receiving waters. Distinct Accumulibacter clades often co-exist in full-scale wastewater treatment plants and lab-scale enrichment bioreactors, and have been hypothesized to inhabit distinct ecological niches. However, since individual strains of the Accumulibacter lineage have not been isolated in pure culture to date, these predictions have been made solely on genome-based comparisons and enrichments with varying strain composition. Here, we used genome-resolved metagenomics and metatranscriptomics to explore the activity of co-existing Accumulibacter strains in an engineered bioreactor environment. We obtained four high-quality genomes of Accumulibacter strains that were present in the bioreactor ecosystem, one of which is a completely contiguous draft genome scaffolded with long Nanopore reads. We identified core and accessory genes to investigate how gene expression patterns differed among the dominating strains. Using this approach, we were able to identify putative pathways and functions that may confer distinct functions to Accumulibacter strains and provide key functional insights into this biotechnologically significant microbial lineage.

**IMPORTANCE:** ‘*Candidatus* Accumulibacter phosphatis’ is a model polyphosphate accumulating organism that has been studied using genome-resolved metagenomics, metatranscriptomics, and metaproteomics to understand the EBPR process. Within the Accumulibacter lineage, several similar but diverging clades are defined by shared sequence identity of the polyphosphate kinase (*ppk1)* locus. These clades are predicted to have key functional differences in acetate uptake rates, phage defense mechanisms, and nitrogen cycling capabilities. However, such hypotheses have largely been made based on gene-content comparisons of sequenced Accumulibacter genomes, some of which were obtained from different systems. Here, we performed time-series genome-resolved metatranscriptomics to explore gene expression patterns of co-existing Accumulibacter clades in the same bioreactor ecosystem. Our work provides an approach for elucidating ecologically relevant functions based on gene expression patterns between closely related microbial populations.

## INTRODUCTION

Naturally occurring microbial assemblages are composed of closely related species or subpopulations and harbor extensive genetic diversity. Coherent lineages are often comprised of distinct clades, ecotypes, or “backbone subpopulations” defined by high within-group sequence similarities that translate to ecologically relevant diversity (1–3). For example, genetically diverse ecotypes of the marine cyanobacterium *Prochlorococcus* that are 97% identical by their 16S ribosomal RNA gene sequences exhibit markedly different light-dependent physiologies and distinct seasonal and geographical patterns (1, 2, 4, 5). Closely related *Nitrospira* representing species-like lineages that coexisted for long periods of time in a full-scale wastewater treatment plant exhibited functional differences in preferences for nitrite concentrations and carbon substrates (6). However, dissecting the emergent ecological properties of species-like coherent microbial lineages is still a grand challenge in microbial ecology given that these principles are likely not universal among the breadth of microbial diversity, and is further hindered when there is a lack of pure cultures (7–9).

Enhanced Biological Phosphorus Removal (EBPR) is an economically and environmentally significant process for removing excess phosphorus from wastewater. This process depends on polyphosphate accumulating organisms (PAOs) that polymerize inorganic phosphate into intracellular polyphosphate. ‘*Candidatus* Accumulibacter phosphatis’ (hereafter referred to as Accumulibacter) is a member of the *Betaproteobacteria* in the *Rhodocyclaceae* family and has long been a model PAO (10, 11). Accumulibacter achieves net phosphorus removal through cyclical anaerobic-aerobic phases of feast and famine. In the initial anaerobic phase, abundantly available volatile fatty acids (VFAs) such as acetate are converted to polyhydroxyalkanoates (PHAs), a carbon storage polymer. However, PHA formation requires significant ATP input and reducing power, which is supplied through polyphosphate and glycogen degradation. In the subsequent aerobic phase in which oxygen is available as a terminal electron acceptor and soluble carbon sources are depleted, Accumulibacter uses PHA reserves for growth. Accumulibacter recovers energy during aerobic PHA degradation by replenishing polyphosphate and glycogen reserves, which can be later used for future PHA formation as the anaerobic/aerobic cycle repeats. This feast-famine oscillation through sequential anaerobic-aerobic cycles operating on minute-to-hourly time scales gives rise to net phosphorus removal and the overall EBPR process (10, 12). A few ‘omics-based studies have revealed potential regulatory modules that could coordinate Accumulibacter’s complex physiological behavior during these very dynamic environmental conditions (13–16), but few have attempted to assign unique gene expression dynamics to specific clades (17).

Accumulibacter can be subdivided into two main types (I and II) and further subdivided into multiple clades based upon polyphosphate kinase (*ppk1*) sequence identity since the 16S rRNA marker is too highly conserved among the breadth of known Accumulibacter lineages to resolve species-level differentiation (18–21). The two main types share approximately 85% nucleotide identity across the *ppk1* locus (21) which mirrors genome-wide average nucleotide identity (ANI) (22). The persistence of several phylogenetically distinct clades within the Accumulibacter lineage in both full-scale wastewater treatment plants and lab-scale enrichment bioreactors suggests that different clades may inhabit distinct ecological niches (17, 23). This is largely supported by comparative genomics of Accumulibacter metagenome-assembled genomes (MAGs) recovered from different bioreactor systems and classified using the *ppk1* locus (14, 17, 22, 24–27). Members of different clades have been hypothesized to vary in their acetate uptake rates, nitrate reduction capacity, and phage defense mechanisms (22, 26). For example, clade IA seems to exhibit higher acetate uptake rates and phosphate release rates, whereas evidence exists that clade IIA can reduce nitrate but clade IA cannot (25, 28). However, the full breadth of ecological differentiation among Accumulibacter clades is not yet clear.

Here, we used genome-resolved metagenomics and time-series metatranscriptomics to investigate gene expression patterns of co-existing Accumulibacter strains (representing two main clades) during a standard EBPR cycle. We assembled four high-quality genomes of the dominant (clades IA and IIC) and low abundant (clades IIA and IIF) Accumulibacter strains present in the bioreactor ecosystem. Additionally, we constructed a contiguous draft genome of an Accumulibacter clade IIC genome achieved using long Oxford Nanopore Technology (ONT) reads, representing one of the highest-quality genomes for this clade recovered to date. Using these assembled genomes, we performed time-series RNA-sequencing throughout a typical EBPR feast-famine cycle to study expression profiles of different Accumulibacter strains, with an emphasis on core and flexible gene content. More generally, our work reveals putative ecological functions of co-existing taxa within a dynamic ecosystem.

## MATERIALS AND METHODS

### Bioreactor Operation and Sample Collection

A laboratory-scale sequence batch reactor (SBR) was seeded with activated sludge from the Nine Springs Wastewater Treatment Plant (Madison WI, USA) during August 2015. The bioreactor was operated to simulate EBPR as described previously (24). Briefly, an SBR with a 2-L working volume was operated with the established biphasic feast-famine conditions: a 6-h cycle consisted of the anaerobic phase (sparging with N_2_ gas), 110-min; aerobic (sparging with oxygen), 180-min; settle, 30-min; draw and feed with an acetate-containing synthetic wastewater feed solution, 40-min. A reactor hydraulic residence time of 12-h was maintained by withdrawing 50% of the reactor contents after the settling phase, then filling the reactor with fresh nutrient feed; a mean solids retention time of 4-d was maintained by withdrawing 25% of the mixed reactor contents each day immediately prior to a settling phase. Allylthiourea was added to the synthetic feed solution to inhibit ammonia oxidation and indirectly inhibit nitrification/denitrification.

Three biomass samples for metagenomic sequencing were collected in September of 2015 by centrifuging 2-mL of mixed liquor at 8,000 × *g* for 2 min, and DNA was extracted using a phenol:chloroform extraction protocol. Seven samples for RNA sequencing were collected throughout a single EBPR cycle (3 in the anaerobic phase, 4 in the aerobic phase) in October of 2016 following the sampling strategy conducted by Oyserman et al. (13). Samples were flash-frozen on liquid nitrogen immediately after centrifuging and discarding the supernatant. RNA was extracted using a TRIzol-based extraction method (Thermo Fisher Scientific, Waltham, MA) followed by phenol-chloroform separation and RNA precipitation. RNA was purified following an on-column DNAse digestion using the RNase-Free DNase Set (Qiagen, Venlo, Netherlands) and cleaned up with the RNeasy Mini Kit (Qiagen, Venlo, Netherlands). Accumulibacter clade quantification was performed on biomass samples collected on the same day of the metatranscriptomics experiment using clade-specific *ppk1* qPCR primers as described by Camejo et al. (23).

### Metagenomic and Metatranscriptomic Library Construction and Sequencing

Metagenomic libraries were prepared by shearing 100 ng of DNA to 550 bp long products using the Covaris LE220 (Covaris) and size selected with SPRI beads (Beckman Coulter). The fragments were ligated with end repair, A-tailing, Illumina compatible adapters (IDT Inc) using the KAPA-Illumina library preparation kit (KAPA Biosystems). Libraries were quantified using the KAPA Biosystem next-generation sequencing library qPCR kit and ran on a Roche LightCycler 480 real-time PCR instrument. The quantified libraries were prepared for sequencing on the Illumina HiSeq platform using the v4 TruSeq paired-end cluster kit and the Illumina cBot instrument to create a clustered flow cell for sequencing. Shotgun sequencing was performed at the University of Wisconsin – Madison Biotechnology Center DNA Sequencing facility on the Illumina HiSeq 2500 platform with the TruSeq SBS sequencing kits, followed by 2×150 indexing. Raw metagenomic data consisted of 219.4 million 300-bp Illumina HiSeq reads with approximately 3.9 Gbp per sample. Nanopore sequencing was performed according to the user’s manual version SQK-LSK108.

Total RNA submitted to the University of Wisconsin-Madison Biotechnology Center was verified for purity and integrity using a NanoDrop 2000 Spectrophotometer and an Agilent 2100 BioAnalyzer, respectively. RNA-Seq paired-end libraries were prepared using the TruSeq RNA Library Prep Kit v2 (Illumina, San Diego, CA). Each sample was processed for ribosomal depletion, using the Ribo-Zero™ rRNA Removal Kit (Bacteria). mRNA was purified from total RNA using paramagnetic beads (Agencourt RNAClean XP Beads, Beckman Coulter Inc., Brea, CA) and fragmented by heating in the presence of a divalent cation. The fragmented RNA was then converted to cDNA with reverse transcriptase using SuperScript II Reverse Transcriptase (Invitrogen, Carlsbad, California, USA) with random hexamer priming and the resultant double-stranded cDNA was purified. cDNA ends were repaired, adenylated at the 3’ ends, and then ligated to Illumina adapter sequences. The quality and quantity of the DNA were assessed using an Agilent DNA 1000 series chip assay and Thermo Fisher Qubit dsDNA HS Assay Kit. Libraries were diluted to 2nM, pooled in an equimolar ratio, and sequenced on an Illumina HiSeq 2500, using a single lane of paired-end, 100 bp sequencing, and v2 Rapid SBS chemistry.

### Metagenomic Assembly and Annotation

We applied two different assembly and binning approaches to obtain high-quality genomes of the dominant Accumulibacter strains present in the reactors. For the first approach, Illumina unmerged reads were quality filtered and trimmed using the Sickle software v1.33 (29). Reads were merged with FLASH v1.0.3 22 (30), with a mismatch value of ≤0.25 and a minimum of ten overlapping bases from paired sequences, resulting in merged read lengths of 150 to 290 bp. FASTQ files were then converted to FASTA format using the Seqtk software v1.0 (31). Metagenomic reads from all three samples were then co-assembled using the Velvet assembler with a k-mer size of 65 bp, a minimum contig length of 200 bp, and a paired-end insert size of 300 bp (32). Metavelvet was used to improve the assembly generated by Velvet (33). Metagenomic contigs were then binned using Maxbin (34). This approach allowed us to assemble high-quality genomes of clades IIA and IIC, named UW5 and UW6, respectively. Long Nanopore reads were generated under the standard protocol and used to scaffold assembled contigs with MeDuSa (35), and manually inspected to remove short contigs and decontaminated using ProDeGe (36) and Anvi’o (37). Further scaffolding was performed on both of these genomes using Nanopore long reads using LINKS (38) and GapCloser (39).

Because this pipeline did not produce high-quality genomes of clades IA and IIF, we applied a different approach to assemble these genomes. All reads from each metagenomic sample were individually and co-assembled assembled using metaSPAdes (40). Reads from each metagenome were mapped against each of the assemblies using bbmap with a 95% sequence identity cutoff to obtain differential coverage (41), and contigs were binned using MetaBat (42). Identical clusters of bins were dereplicated across assemblies using dRep (43) to obtain the best quality genome of both clades IA and IIF, named UW4 and UW7, respectively. All genomes were quality checked using CheckM (44) and functional annotations assigned with Prokka and KofamKOALA (45, 46). We were unable to scaffold the genomes of UW4-IA and UW7-IIF with long Nanopore reads because of their low abundance at the time of metagenomic sequencing.

Each genome was assigned to a particular clade both by comparing the *ppk1* sequence identity and ANI to previously published Accumulibacter genomes. A *ppk1* database was created using sequences from previously published Accumulibacter references and clone sequences. We searched for the *ppk1* gene in our draft genomes using this database and aligned the corresponding hits with MAFFT (47). A phylogenetic tree of aligned *ppk1* gene sequences from Accumulibacter references, select clone sequences, and the outgroups *Dechloromonas aromatica RCB* and *Rhodoc*yclus *tenuis DSM 110* with RAxML v8.1 with 100 rapid bootstraps (48). To use any given Accumulibacter reference genome in downstream comparisons, a confident *ppk1* hit had to be identified using the above methods. As a technical note, the IA-UW2 strain assembled by Flowers et al. (22) was renamed to UW3, as the original assembly contained an additional contig that was likely from a prophage. The prophage contig was removed from the IA-UW2 strain and renamed to UW3, and therefore the UW numerical nomenclature follows in sequential order after UW3. A phylogenetic tree of *ppk1* nucleotide sequences was performed and compared to select clone sequences. Pairwise genome-wide ANI was calculated between the four Accumulibacter draft genomes and all available Accumulibacter genomes in Genbank with FastANI (49). A species tree of all Accumulibacter references and outgroup genomes was constructed using single-copy markers from the GTDBK-tk (50), aligned with MAFFT (47), built with RAxML with 100 rapid bootstraps (48), and visualized in iTOL (51). To compare genome-wide ANI scores of Accumulibacter references with pairwise nucleotide identity of the *ppk1* locus of all references, we performed pairwise BLAST for all *ppk1* coding regions and reported the percent identity (52).

### Identification of Core and Accessory Gene Content

We identified core and accessory gene content of Accumulibacter clades through clustering orthologous groups of genes (COGs) using PyParanoid (53). We used the four Accumulibacter MAGs generated in this study (Table 1) and high-quality Accumulibacter reference genomes (above 90% completeness and less than 5% redundancy) for this analysis, with *Dechloromonas aromatica RCB* and *Rhodocyclus tenuis DSM 110* as outgroups (Accumulibacter references listed in Supplementary Table 1 available at https://figshare.com/articles/dataset/Accumulibacter_References_Metadata/13237148). Briefly, an all-vs-all comparison of proteins for all genomes was performed using DIAMOND (54), pairwise homology scores calculated with the InParanoid algorithm (55), gene families constructed with MCL (56), and HMMs built for each gene family with HMMER (57). A phylogenetic tree was constructed for genomes used for the COGs analysis by using the coding regions of the *ppk1* locus and overlaid with the presence/absence of core and flexible gene content using the ete3 python package (58). A similarity score between UW Accumulibacter genomes was calculated as the pairwise Pearson correlation coefficient between gene families using the numpy python package (59). The presence and absence of different gene families were visualized in an upset plot between UW Accumulibacter genomes using the ComplexUpset package based on the UpSetR package in R (60). All data wrangling and plotting were performed using the tidyverse suite of packages in R (61).

**Table 1.** Assembled Accumulibacter Genome Statistics. Genome quality calculations for completeness and contaminations were made with CheckM based on the presence/absence of single-copy genes (44).

| **Genome** | **Completion** | **Contamination** | **Size_Mbp** | **Contigs** | **GC** |
| --- | --- | --- | --- | --- | --- |
| IIA-UW5 | 98.99 | 5.24 | 4.88 | 68 | 64.3 |
| IIC-UW6 | 98.57 | 2.46 | 5.18 | 1 | 61.1 |
| IIF-UW7 | 98.97 | 3.04 | 4.87 | 102 | 66.3 |
| IA-UW4 | 92.90 | 3.37 | 4.29 | 358 | 64.0 |

### Metatranscriptomics Mapping and Processing

Metatranscriptomics reads from each of the seven samples were quality filtered using fastp (62) and rRNA removed with SortMeRNA (63). The coding regions of all four Accumulibacter genomes assembled in this study (Table 1) were predicted with Prodigal (64) and annotated with both Prokka and KofamKOALA (45, 46), and concatenated together to create a mapping index. Metatranscriptomic reads were competitively pseudoaligned to the reference index and counts quantified with kallisto (65). Genes with a sum of more than 10,000 counts across all 7 samples were removed, as these are likely ribosomal RNAs that were not removed previously. Additionally, genes annotated as rRNAs by barrnap were manually removed (45).

To explore genes and metabolic functions that may exhibit hallmark anaerobic-aerobic feast-famine cycling patterns among the dominant IA-UW4 and IIC-UW6 strains, we identified sets of genes that were differentially expressed between anaerobic and aerobic conditions using DESeq2 (66). Differential gene expression calculations were performed for each strain separately to account for differences in the relative abundance of metatranscriptomic reads mapping back to each of the four MAGs. Low-count genes were removed individually for each genome by requiring that all genes had to have more than 10 counts mapped across 3 or more samples. For IA-UW4, 3549 genes of a total 3919 predicted genes remained after applying this filter, and for IIC-UW6, 4297 genes of a total 4877 predicted genes remained after applying this filter. We analyzed expression patterns of differentially expressed genes using a threshold cutoff of ±1.5 log2 fold-change between anaerobic and aerobic conditions and then identified differentially expressed genes as core or accessory among clades IA (str. UW4) and IIC (str. UW6). For the core group analysis, the corresponding ortholog in each genome had to meet the minimum count threshold as described above to include in the comparison, with genes in one or both of the strains being differentially expressed. For the accessory analysis for each strain, the gene had to be differentially expressed and not contained in the other genome but could be present in other strains or other genomes belonging to that clade.

## Data and Code Availability

Raw sequencing files for the three metagenomes and seven metatranscriptomes and genome assemblies for the four Accumulibacter genomes are available at NCBI under the BioProject accession PRJNA668760. All genome assemblies used in this study in their current forms as well as transcriptomic count tables are available at: https://figshare.com/projects/Metabolic_Plasticity_of_Accumulibacter_Clades/90614. All code is available at https://github.com/elizabethmcd/R3R4.

## RESULTS

### Enrichment of Co-Existing Accumulibacter Clades

We inoculated a 2-L sequencing-batch reactor with activated sludge from a full-scale wastewater treatment plant in Madison, WI, USA and maintained operation for 40 months. The reactor was primarily fed with acetate and operated under cyclic anaerobic-aerobic phases with a 4-day solids retention time to simulate the EBPR process and enrich for Accumulibacter (Figure 1). During a period of stable EBPR (14 months following inoculation) in which acetate was completely diminished by the start of the aerobic phase and soluble phosphate was below 1.0 mg-L^-1^ at the end of the aerobic phase, we collected seven samples across a single cycle (three in the anaerobic phase, four in the aerobic phase) for RNA-sequencing (Figure 1A). We performed qPCR of the *ppk1* as described previously (23) to quantify Accumulibacter clades present in the reactor on the same day as the RNA-seq experiment (Figure 1B). At the time of the RNA-seq experiment, the bioreactor was primarily enriched in Accumulibacter clade IA, followed by clade IIC and clade IIA.

**Figure 1.**
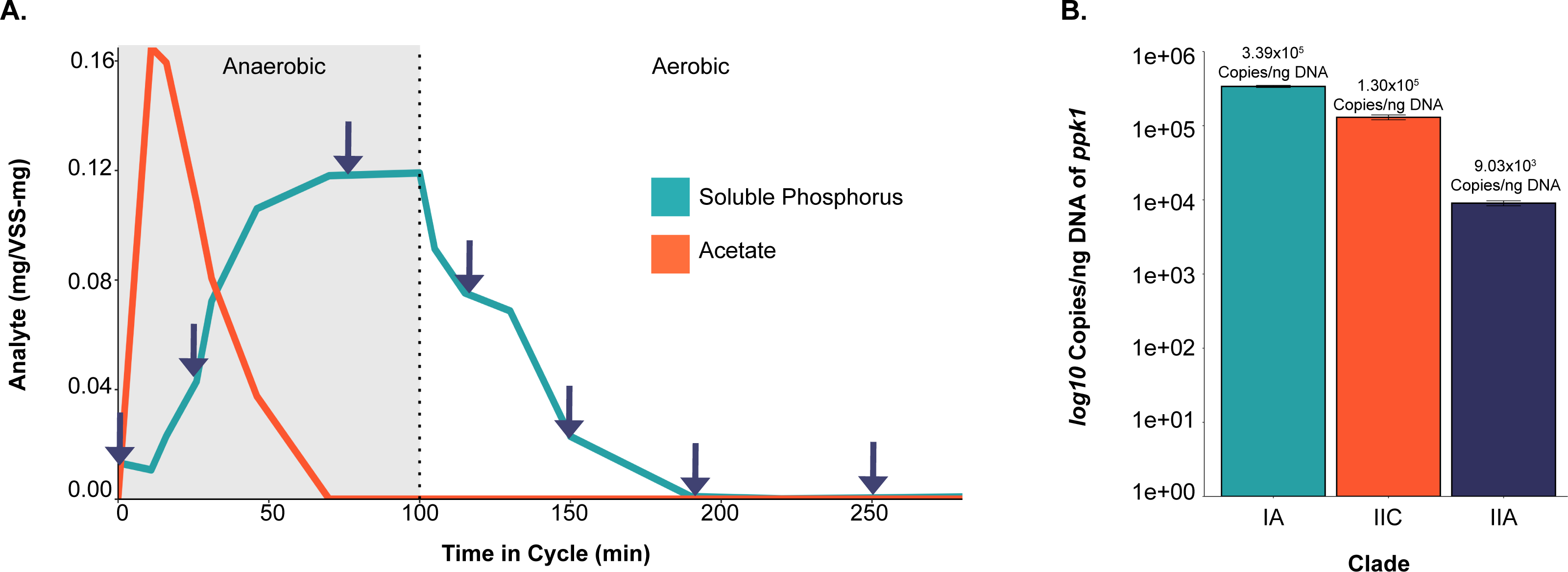
Overview of Experimental Design and Enriched Accumulibacter Clades. **1A.** Overview of EBPR cycle and sampling scheme. Samples were collected for RNA sequencing over the course of a normal EBPR cycle, with reported measurements for soluble phosphorus and acetate normalized by volatile suspended solids (VSS). Samples were taken at time-points 0, 31, 70, 115, 150, 190, and 250 minutes. **1B.** Abundance of different Accumulibacter clades by *ppk1* copies per ng of DNA. Quantification of Accumulibacter clades was performed by qPCR of the *ppk1* gene as described in Camejo et al. (23).

From metagenome samples collected from this bioreactor earlier, we assembled high-quality genomes of belonging to four different Accumulibacter clades (Table 1), including the dominant IA and IIC (strains IA-UW4 and IIC-UW6, respectively) clades and the less abundant IIA and IIF clades (strains IIA-UW5 and IIF-UW7, respectively) (Figure 2). Each of the assembled genomes falls within the established Accumulibacter clade nomenclature as defined by *ppk1* sequence identity and phylogenetic placement when compared to other publicly available Accumulibacter genomes and clone sequences (Figure 2A). Additionally, the *ppk1* hierarchical structure is also reflected by pairwise-average nucleotide identity (ANI) boundaries between clades, with some exceptions for newly assembled Accumulibacter genomes that fall outside of the established ‘*C.* Accumulibacter phosphatis’ lineage (Figure 2B). All assembled genomes are above 90% complete and contain less than 5% redundancy as calculated by CheckM (44) (Table 1). We were able to scaffold the assemblies of strains IIA-UW5 and IIC-UW6 using long Nanopore reads, constructing a completely contiguous draft genome of the IIC-UW6 strain. To our knowledge, the assembled IIC-UW6 genome is the most contiguous, highest-quality reference genome available for this clade, providing a valuable new Accumulibacter reference genome (Figure 2C).

**Figure 2.**
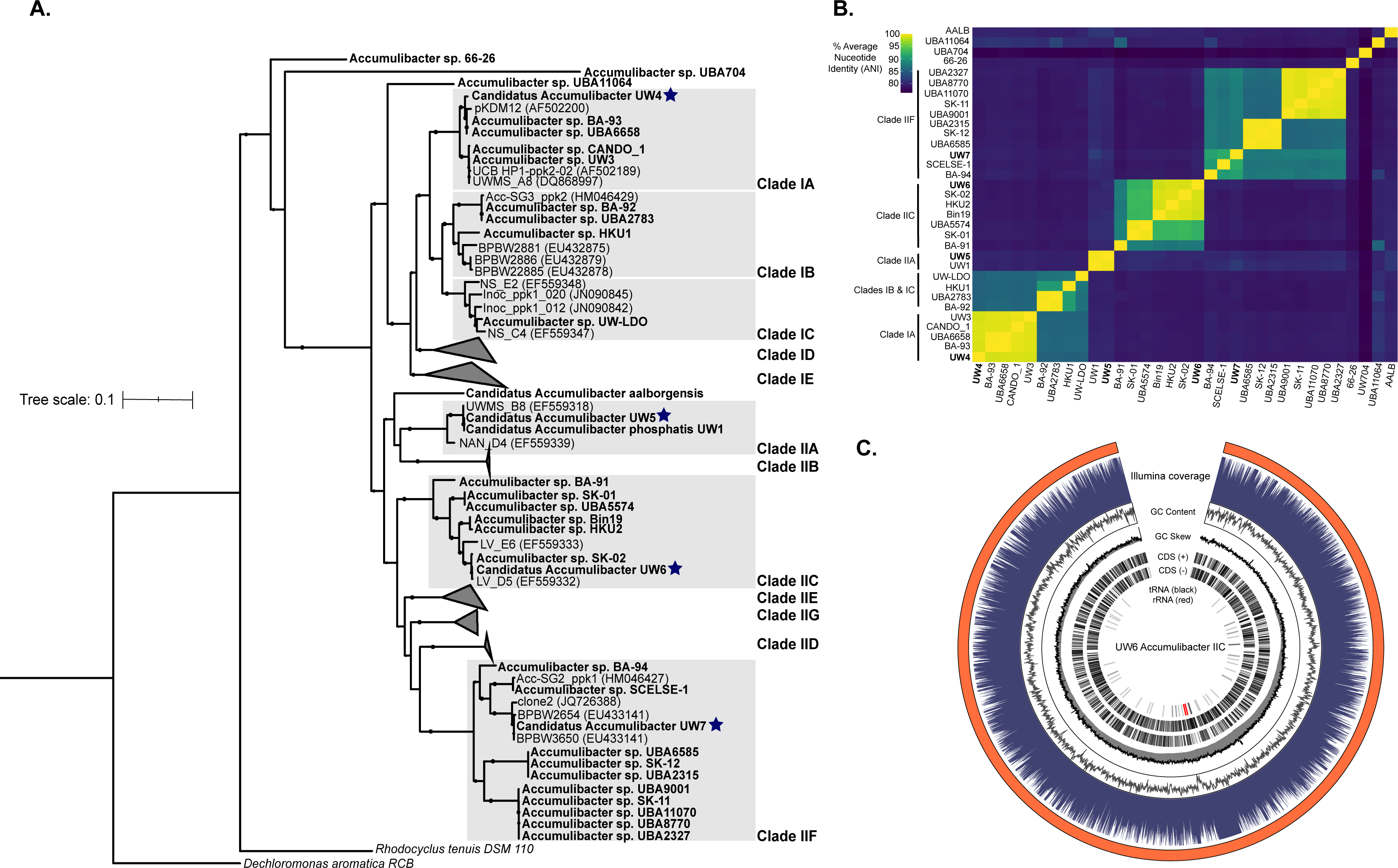
Assembly of Four High-Quality Accumulibacter Genomes. **2A.** Phylogenetic tree of *ppk1* nucleotide sequences from available Accumulibacter reference genomes, clone sequences, and the four assembled Accumulibacter genomes from this study. Tips in bold represent *ppk1* sequences from metagenome-assembled genomes, whereas others are sequences from clone fragments available at the referenced accession numbers. Starred tips represent *ppk1* sequences from the four assembled Accumulibacter genomes from this study. Tree was constructed using RaXML with 100 rapid bootstraps. **2B.** Pairwise genome-wide ANI comparisons of all publicly available Accumulibacter references and the four new Accumulibacter genomes, denoted in bold. Genomes are grouped according to clade designation by *ppk1* sequence identity and in order of the *ppk1* phylogeny, except for those that do not fall in an established clade. **2C.** Genome diagram of the UW6 Accumulibacter IIC MAG (98.6% completeness, 2.5% contamination, 5.18 Mbp). Layers represent from top to bottom: coverage over 1000 bp windows, GC content and skew across the same 1000 bp windows, coding sequences on the positive and negative strands, and positions of predicted rRNA (red) and tRNA (black) sequences.

### Distribution of Shared and Flexible Gene Content between Clades

We characterized the functional diversity of different Accumulibacter clades through clustering orthologous groups of genes (COGs) (Figure 3A). We repeated a COGs analysis from Skennerton et al. (26) and Oyserman et al. (67), but included an updated set of high-quality Accumulibacter references, the UW-generated genomes, and the outgroups *Dechloromonas aromatica* and *Rhodocyclus tenuis* (Figure 3). The Accumulibacter lineage exhibits a tremendous amount of functional diversity, with only approximately 25% of COGs shared amongst all Accumulibacter clades (Figure 3A). Additionally, there is extensive diversity within the two Accumulibacter types and within individual clades, with only a few representatives of each clade more than approximately 75% similar to each other by gene content (Supplementary Figure 1). Specifically, clade IIC broadly harbors the most diversity of all sampled genomes by gene content, with representatives of this clade not sharing more than 25% of COGs (Figure 3A). Other clades are more similar by gene content, but this could be due to fewer genomes sampled from these clades. Most of the highest-quality, publicly available Accumulibacter genome references belong to clades IIF and IIC.

**Figure 3.**
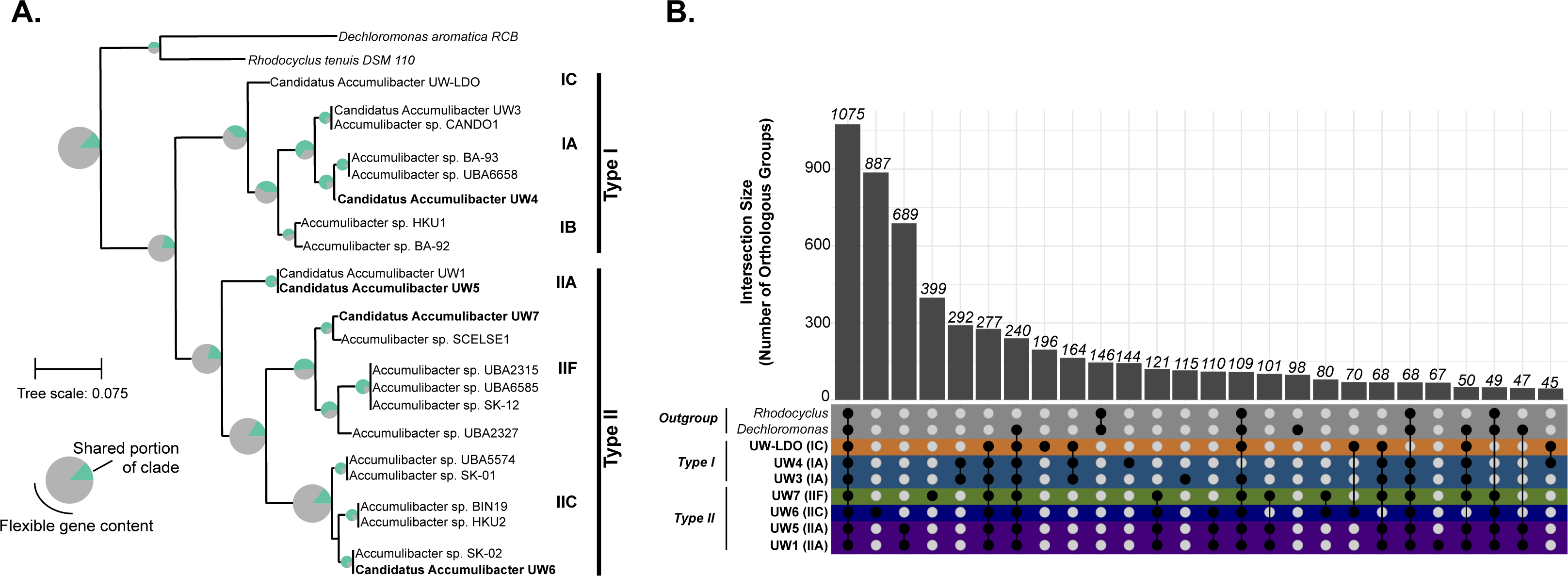
Core and Flexible Gene Content among the Accumulibacter lineage and UW genomes. **3A.** Phylogenetic tree of the coding regions for the *ppk1* locus among high-quality Accumulibacter references, UW genomes, and outgroup taxa *Dechloromonas aromatica RCB* and *Rhodocyclus tenuis DSM 110.* The pie graph at each node represents the fraction of genes that are shared and flexible for genomes belonging to that clade. **3B.** Upset plot representing the intersections of groups of orthologous gene clusters among UW genomes and outgroups, analogous to a Venn diagram. 1075 gene families were shared among all UW Accumulibacter clades and outgroup genomes and 277 separate gene families were found only within the UW Accumulibacter clades. Each bar represents the number of orthologous gene clusters, and the dot plot represents the genomes in which the groups intersect.

We then compared COGs between Accumulibacter clades collected from bioreactors seeded from the Nine Springs Wastewater Treatment Plant in Madison, as these genomes represent a shared origin (Figure 3B). In addition to two lab-scale bioreactors harboring distinct Accumulibacter clades (strains UW1 and UW3, and UW4-7, respectively) (22, 24), we recently characterized clade IC (strain UW-LDO-IC) using genome-resolved metagenomics and metatranscriptomics (14). Overall, genomes in Type I and Type II cluster respectively, where genomes within Type I are overall more similar to each other, as expected (Supplementary Figure 1). Clade IIA genomes (strains UW1 and UW5) were assembled from separate enrichments derived from the same full-scale treatment plant approximately 12 years apart, but are very similar by COG presence/absence (Supplementary Figure 1). UW genomes within Type II are less similar by gene content and seem to be more diverse in this regard, though again this could be due to under-sampling of Type I (8 genomes in Type I and 14 in Type II). Additionally, clade IIF contains more overlap in gene content with Type I genomes in gene families than some Type II genomes do to each other (Supplementary Figure 1).

COG overlap and uniqueness is shown by the intersection of COG groups between UW Accumulibacter clades and the *Rhodocyclus* and *Decholoromonas* outgroups (Figure 3B). There are 1075 gene families shared among all UW Accumulibacter clades and outgroup genomes, with 277 separate gene families only within the UW Accumulibacter clades (Figure 3B). Among individual clades, there are 887 groups only within clade IIC (UW6), 399 within clade IIA (UW1 and UW5), 399 within IIF (UW7), 292 within clade IA (UW3 and UW4), and 196 within IC (UW-LDO). Although clade IIC str. UW6 contains more overlap in gene content with SK-02 (Figure 3A), the intersection analysis (Figure 3B) identified gene families that are available to individual strains within the same bioreactor ecosystem, highlighting the potential for functional differentiation.

### Transcriptional Profiles of Accumulibacter Strains

Although *ppk1* locus quantification showed that the bioreactor was mostly enriched in clade IA (see Figure 1B), surprisingly, more RNA-seq reads mapped to the clade IIC-UW6 genome than to the clade IA-UW4 (Figure 4A). Approximately 40 million total transcriptional reads aligned to the strain IIC-UW6 genome across the full time-series experiment, whereas strains IA-UW4 and IIA-UW5 recruited roughly the same number of reads, with 14 million and 11 million, respectively (Table 2). Markedly lower read levels mapped to the IIF-UW7 strain relative to the other three clades, and therefore we do not include the IIF-UW7 strain in our subsequent analyses. We explored the top differentially expressed genes in strain IIC-UW6, comparing anaerobic and aerobic conditions (Figure 4B). Genes upregulated in the anaerobic phase belonged to pathways for central-carbon transformations, fatty acid biosynthesis, and PHA synthesis. Interestingly, all genes within the high-affinity phosphate transport system Pst were upregulated in the aerobic phase. This includes the substrate-binding component *pstS, phoU* regulator, and the ABC-type transporter *pstABC* complex (Figure 4B).

**Figure 4.**
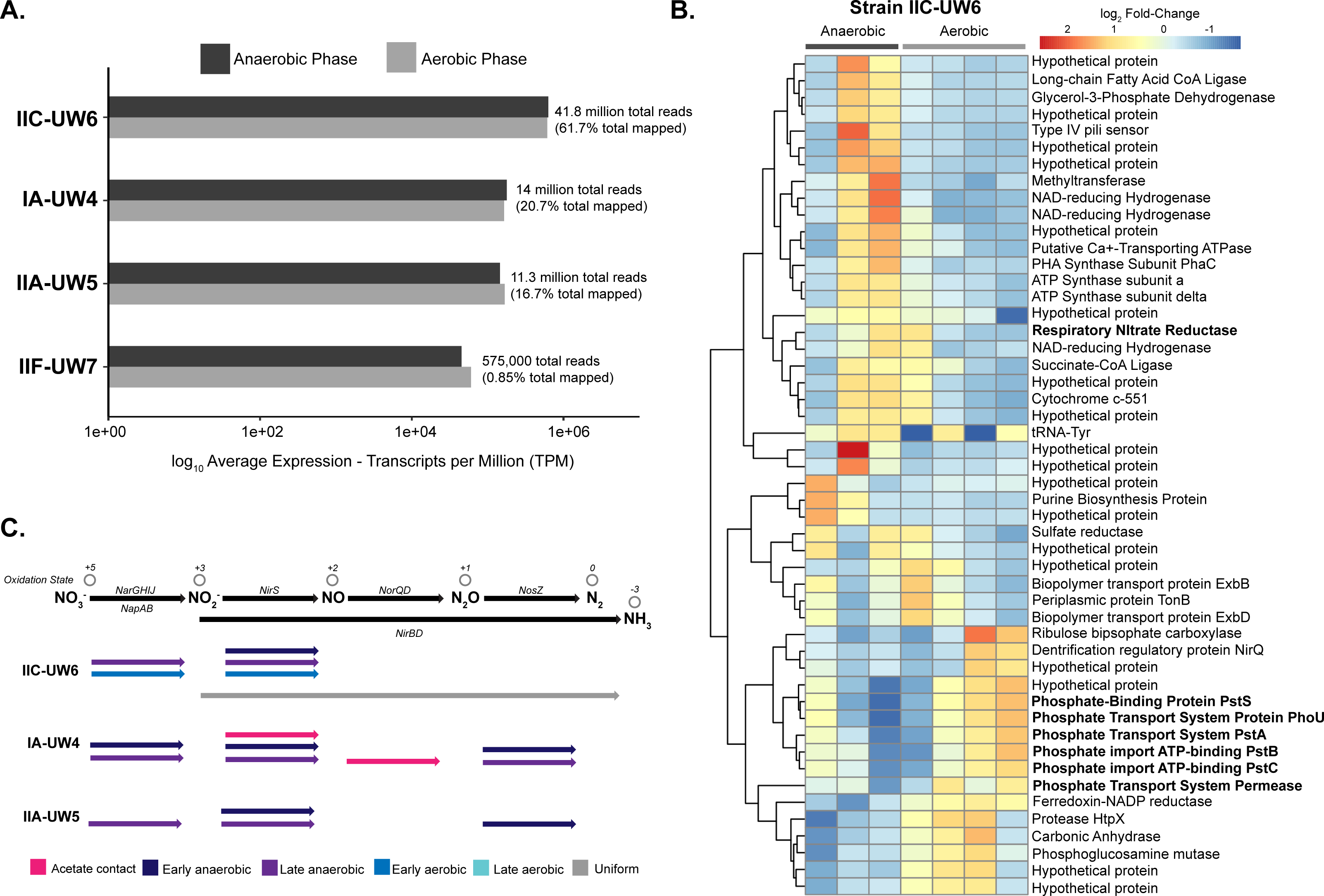
Transcriptomic Profiles of Accumulibacter Clades. **4A.** Average expression profiles of each reference genome in the anaerobic and aerobic phases. The IIC-UW6 genome mapped the most transcripts even though the IA-UW4 was more abundant. Reads were competitively mapped to the four assembled Accumulibacter clade genomes and normalized by transcripts per million (TPM). The sum of the normalized counts of the three samples in the anaerobic phase and four samples in the aerobic phase were averaged, respectively, and plotted on a log_10_ scale. Numbers and percentages for adjacent to normalized bars represent the raw number of reads that are mapped to each genome. **4B.** Top 50 differentially-expressed genes in clade IIC-UW6. All 50 genes pass the threshold of ±1.5 log-10 fold-change across the cycle. Annotations were predicted from a combination of Prokka (45) and KofamKOALA (46). **4C.** Denitrification gene expression among clades IIC-UW6, IA-UW4, and IIA-UW5. We summarized expression for genes involved in denitrification by highlighting parts of the cycle in which certain genes are expressed the most. This is characterized by acetate contact (pink), early anaerobic (dark purple), late anaerobic (purple), early aerobic (dark blue) and late aerobic (light blue), where uniform describes uniform expression across the entire cycle (grey). If an arrow is missing for a step, that particular clade did not contain confident annotations within the genome for those genes. For nitrate reduction, the *nar* system is represented as a black dot and the *nap* system is represented as a grey dot. Multiple arrows are shown for a single step in which particular genes were highly expressed in multiple phases.

**Table 2.**
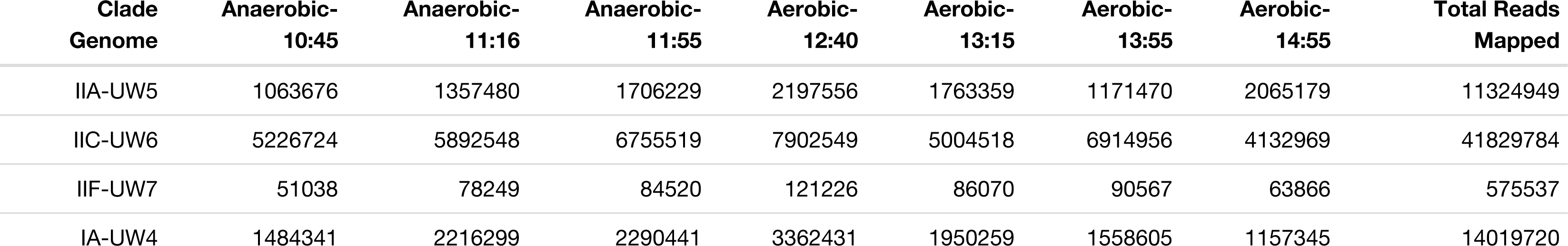
Metatranscriptomic Reads Mapped to Accumullibacter Genomes. Counts of raw metatranscriptomic reads mapped to each Accumulibacter genome with kallisto (65) in each sample for the three anaerobic samples and four aerobic samples and total reads mapped for all samples.

The co-existing clades in the bioreactor system differ in both the presence of sets of nitrogen cycling gene sets and their expression profiles (Figure 4C). In strain IA-UW4, we detected the *napAGH* genes for nitrate reduction, whereas we did not detect *napB*, possibly due to these genes being close to the end of a contig in this genome. We also detected a *nirS*-like nitrite reductase, both subunits of the nitric oxide reductase *norQD,* and the nitrous oxide reductase *nosZ* in strain IA-UW4. Most of these genes in strain IA-UW4 exhibit the highest expression patterns in the early and late anaerobic stages (Figure 4C). Conversely, strain IIC-UW6 contains the *narGHIJ* machinery for nitrate reduction, a different *nirS-*like nitrite reductase than in IA-UW4, and does not contain a nitrous oxide reductase. However, strain IIC-UW6 does contain the nitrite reductase *nirBD* for reducing nitrite to ammonium, which is uniformly expressed across the cycle (Figure 4C).

### Expression Dynamics of Genes Shared by Strains IA-UW4 and IIC-UW6

When characterizing expression dynamics of COGs shared between strains IA-UW4 and IIC-UW6, we found very few orthologs that exceeded the 1.5-fold change cutoff for differential expression in both strains (Figure 5). These genes include a coproporphyrinogen III oxidase, NAD-reducing HoxS subunits, the *phaC* subunit of the PHA synthase, the long-chain fatty-acid-CoA ligase FadD13, and several hypothetical proteins. The fatty-acid CoA-ligase incorporates ATP, CoA, and fatty acids of variable length to form acyl-CoA to be degraded for energy production, incorporated into complex lipids, or used in other metabolic pathways (68). The acyl-CoA ligase is upregulated in the anaerobic phase of both clades, suggesting a role in anaerobic carbon metabolism, as acyl-CoA can ultimately form acetyl-CoA through beta-oxidation. The *phaC* subunit of the PHA synthase is differentially expressed in the anaerobic phase for both clades, as is typical of the hallmark EBPR metabolism for Accumulibacter (13, 69).

**Figure 5.**
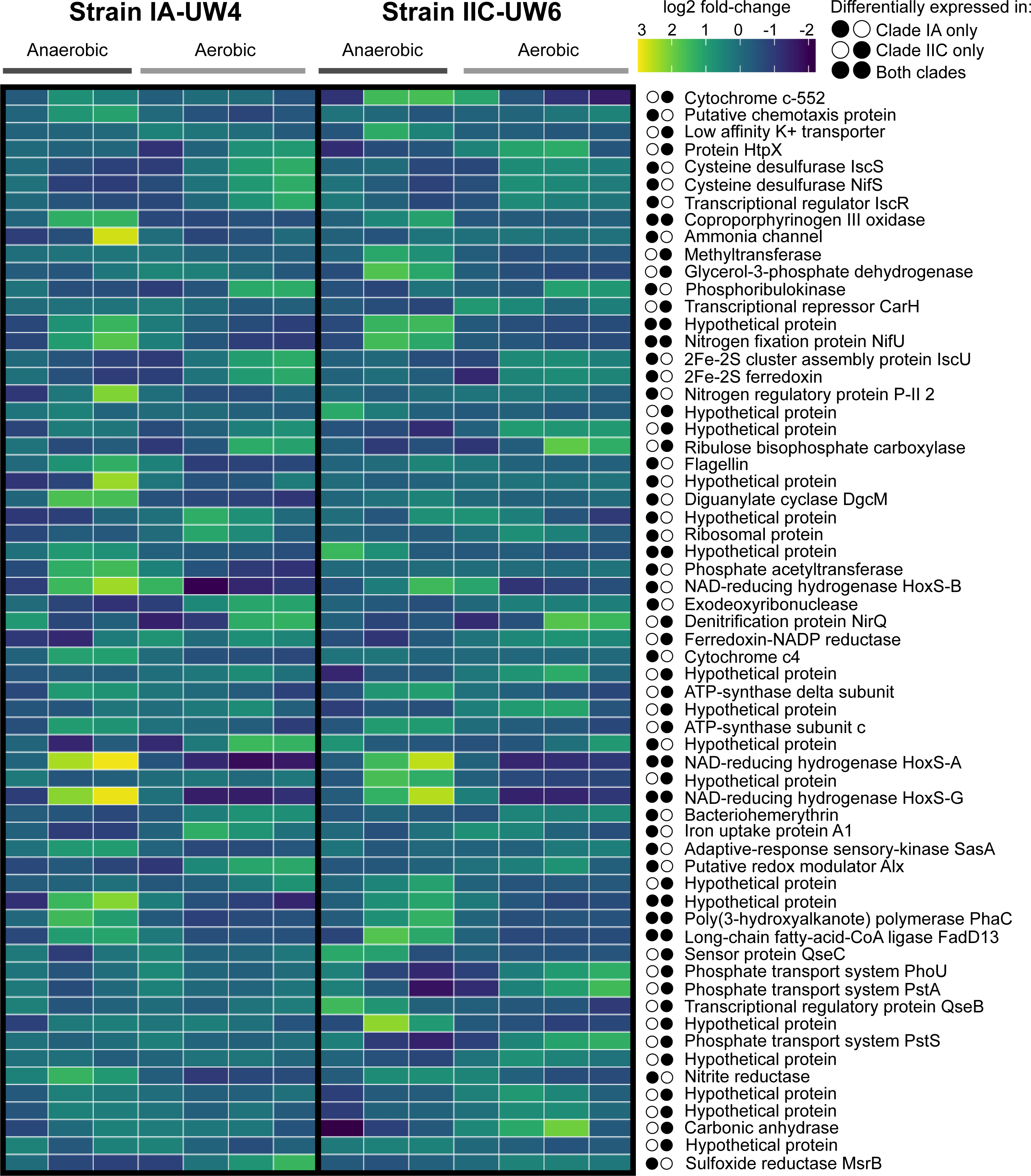
Differential Expression of Shared Genes in Strains IA-UW4 and IIC-UW6. Gene expression profiles of core COGs across the EBPR cycle that are differentially expressed between anaerobic and aerobic conditions in either IA-UW4 or IIC-UW6 or both strains. To consider a gene differentially expressed in a strain, the COG must exhibit ±1.5 fold-change in expression between anaerobic and aerobic phases. Homologs that are directly compared to each other in IA-UW4 and IIC-UW6 are orthologous genes that belong to the same cluster.

Additionally, both the IA-UW4 and IIC-UW6 strains carry most of the subunits for the bidirectional NiFe Hox system, encoding an anaerobic hydrogenase. The HoxH and HoxY beta and delta subunits form the hydrogenase moiety, and the FeS-containing HoxF and HoxU alpha and gamma subunits catalyze NAD(P)H/NAD(P)+ oxidation/reduction coupled to the hydrogenase moiety (70–72). In both clades, the FeS cluster HoxF and HoxU alpha and gamma subunits are strongly differentially expressed in the anaerobic phases, as was observed previously in a genome-resolved metatranscriptomic investigation of strain IIA-UW1 (13). However, the IIC-UW6 genome is missing the HoxY delta subunit of the hydrogenase moiety, and only strain IA-UW4 exhibits differential expression of the HoxH beta subunit in the anaerobic phase. Strain IIC-UW6 does exhibit higher expression of HoxH in the late anaerobic and early aerobic phases but does not meet the threshold cutoff for differential gene expression. A prior study focused on strain IIA-UW1 also demonstrated hydrogenase activity in the anaerobic phase, and then demonstrated hydrogen gas production after acetate addition (13). The hydrogen gas production was hypothesized to replenish NAD+ after glycogen degradation. Subsequent comparative genomics analysis suggested that the Hox system is an ancestral state of Accumulibacter (67). Furthermore, recent metabolic models which integrated hydrogen production under periods of excess acetate or glycogen were better able to capture the diversity of observed EBPR stoichiometry (73). In this ecosystem, both strains may exhibit anaerobic hydrogenase activity and thus produce hydrogen, but due to the missing subunit in the IIC-UW6 genome and lack of differential gene expression, this is uncertain.

Interestingly, only one strain exhibited differential expression patterns of several core ortholog subsets important for EBPR-related functions exhibit. For example, subunits of the high-affinity phosphate transporter Pst system are only differentially expressed in strain IIC-UW6 and not IA-UW4, as observed for strain IIC-UW6 above (see Figure 4B). Additionally, nitrogen cycling genes that contain orthologs in both clades display different expression profiles. A nitrite reductase is only differentially expressed in the anaerobic phase in strain IA-UW4 and the denitrification protein NirQ is differentially expressed in only strain IIC-UW6. Whereas the nitrogen fixation protein subunit NifU is differentially expressed in both strains. To activate acetate to acetyl-CoA, the high-affinity acetyl-CoA synthase (ACS) or low-affinity acetate kinase/phosphotransacetylase (AckA/Pta) pathways can be employed. The strain IA-UW4 assembly is missing an acetate kinase, and phosphate acetyltransferase is only differentially expressed in this strain but not in IIC-UW6. Within strain IIC-UW6, both the acetate kinase and phosphate acetyltransferase genes exhibit relatively low levels of expression over the time-series. Both strains contain several acetyl-CoA synthases that are highly expressed in the anaerobic phase, particularly upon acetate contact, but do not exhibit notable differences in differential expression between the two phases. Previous work based on community proteomics and enzymatic assays have suggested that although both activation pathways are present and expressed in Accumulibacter, the high-affinity pathway is preferentially used (74, 75). Although a phosphate acetyltransferase is differentially expressed in strain IA-UW4, this gene has lower transcripts relative to the acetyl-CoA synthase, providing more evidence that the lineage (specifically both clades IA and IIC) may employ the high-affinity system.

### Differential Expression of Flexible Genes in Strain IA-UW4

We lastly characterized expression dynamics of flexible gene content in strain IA-UW4 (Figure 6). Since a majority of the differentially expressed flexible genes in strain IIC-UW6 were annotated as hypothetical proteins, we focused on putative differentiating features of clade IA-UW4. Flexible gene content for IA-UW4 can be defined as any ortholog that is not present in the IIC-UW6 genome, whether or not that ortholog is only contained within clade IA genomes, Type I as a whole, within other Type II or clade IIC genomes, or the *Dechloromonas* or *Rhodocyclus* outgroups. Thus, the absence of a particular ortholog in the strain IIC-UW6 genome but present in the IA-UW4 genome or any other genomes was inferred as a lack of functional capability of IIC-UW6 in this particular ecosystem.

**Figure 6.**
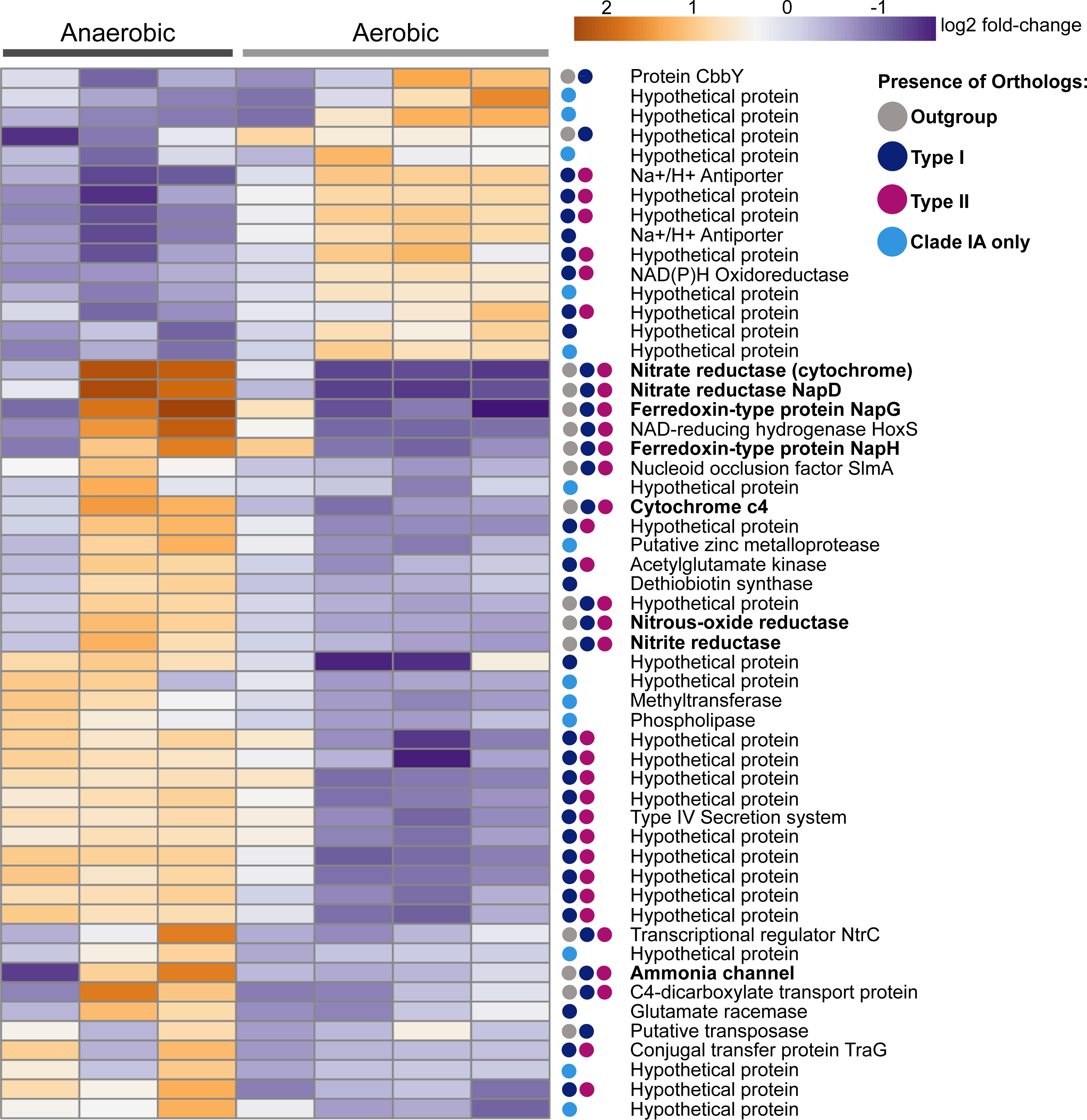
Differential Expression of Accessory Genes in Strain IA-UW4. Gene expression profiles of groups of accessory gene groups in strain IA-UW4 that are not present in the IIC-UW6 genome. A gene was considered differentially expressed between the anaerobic and aerobic conditions if it exhibited greater or less than ±1.5 fold-change in expression. Colored dots represent whether or not that gene contains orthologs within outgroup genomes, other Type II Accumulibacter other than the IIC-UW6 genome, within Type I Accumulibacter, or only within IA.

Particularly striking is the differential expression profiles of nitrogen cycling genes that do not contain orthologs in the IIC-UW6 genome. As stated above, clade IA-UW4 contains the *nap* system whereas strain IIC-UW6 contains the *nar* system for reducing nitrate to nitrite (Figure 4C). In strain IA-UW4, the *napDGH* subunits are differentially expressed between the anaerobic and aerobic phases, and these genes are present in the outgroups, as well as both Accumulibacter types except the IIC-UW6 genome (Figure 6). Both genomes contain a *nirS* nitrite reductase, but the respective homologs are not within the same COG, and thus strain IA-UW4 contains a different *nirS* that is upregulated in the anaerobic phase. Strain IIC-UW6 does not contain both subunits of the nitric-oxide reductase *norQD,* whereas strain IA-UW4 does. These subunits are not differentially expressed above the set threshold but do exhibit the highest expression upon acetate contact in the anaerobic phase (see Figure 4B). Additionally, strain IIC-UW6 does not contain the nitrous-oxide reductase *nosZ* (see Figure 4B), whereas strain IA-UW4 does, and this is differentially expressed in the anaerobic phase (Figure 6). These results suggest that strain IA-UW4 may have the capability for full denitrification from nitrate to nitrogen gas, or at least use the reduction of nitrate/nitrite as a source of energy in the anaerobic phase in a denitrifying bioreactor. Although strain IIC-UW6 does contain the *nirBD* nitrite reductase complex for reducing nitrite to ammonium, these genes are uniformly expressed throughout the cycle (see Figure 4B).

## DISCUSSION

In this work, we performed time-series transcriptomics to explore gene expression profiles of Accumulibacter strains belonging to clades IA and IIC, which co-existed within the same bioreactor ecosystem. By applying genome-resolved metagenomic and metatranscriptomic techniques, we were able to postulate niche-differentiating features of distinct Accumulibacter clades based on core and accessory gene content and differential expression profiles. We assembled high-quality genomes from four Accumulibacter clades and used long reads to generate a single contiguous scaffold of the IIC-UW6 genome. By performing an updated clustering of orthologous groups of genes analysis, we were able to better demonstrate the similarity and uniqueness among Accumulibacter strains from this system. Unexpectedly, although clade IA dominated in abundance by *ppk1* locus quantification, clade IIC was more transcriptionally active. By comparing the expression profiles of COGs shared between strains IA-UW4 and IIC-UW6, we found that genes involved in EBPR-related pathways may differ between the strains with respect to transcription-level regulation. Notably, strain IIC-UW6 showed a strong upregulation of the high-affinity Pst phosphorus transport system in the aerobic phase. High transcription of the Pst system in IIC-UW6 towards the end of the aerobic phase when phosphorus concentrations were lower suggests that the IIC-UW6 strain was more sensitive to bulk phosphate concentrations. This would be congruent with specialization to relatively low phosphorus conditions. Whereas lack of differential expression of the Pst system in IA-UW4 suggests this strain is more specialized under high phosphorus conditions. Strain IA-UW4 contains most of the subunits for the individual key steps for denitrification, and exhibits strong anaerobic upregulation of these genes, suggesting a differentiating role for strain IA-UW4 in nitrogen cycling under denitrifying conditions. Additionally, the IA-UW4 strain contains all subunits for encoding the Hox anaerobic hydrogenase system, which exhibits upregulated gene expression in the anaerobic phase. However, the IIC-UW6 strain is missing one of the subunits even though the other subunits were expressed (but not differentially). Production of hydrogen gas could be a differentiating role for the IA-UW4 strain, as this would impact potential interactions with diverse flanking community members in addition to replenishing NAD+ (13, 24, 76).

The Accumulibacter lineage harbors extensive diversity between and amongst individual clades which is exhibited not only by *ppk1-*based phylogenies, but also by pairwise genome-wide ANI and COG comparisons. For example, strain IIA-UW1 and strain IA-UW3 were present in the same bioreactor ecosystem and only share 85% ANI (22). In the bioreactor enrichment from this study, the dominant strains IA-UW4 and IIC-UW6 share less than 80% ANI (Figure 2B). Recently, a cutoff of >95% ANI has been suggested to delineate distinct bacterial species or cohesive genetic units (49), which is based on observations of coverage gaps at 90-95% sequence identity when metagenomic reads were mapped back to assembled MAGs (77–80). These sequence cutoffs have also been benchmarked against homologous recombination rates and ratios of nonsynonymous to synonymous (*dN/dS*) nucleotide differences (81). Among Accumulibacter genomes adhering to the established *ppk1*-based phylogeny, genomes within individual clades fall within the >95% ANI threshold, whereas ANI comparisons within and between Types I and II are more variable (Figure 2B). Genomes within Type I are similar by approximately 90% ANI, whereas genomes within Type II are not as coherent.

The apparent coexistence of diverse Accumulibacter clades within an ecosystem then raises questions of ecological roles, putative interactions, and evolutionary dynamics of this lineage over time and space. The ecotype model of speciation suggests that cohesive, ecologically distinct ecotypes can coexist because they occupy separate niche spaces (3, 82). Because Accumulibacter has not been isolated in pure culture to date, most ecological and evolutionary inferences have been made from genome-based comparisons (26, 67) or mapping metagenomic reads back to assembled genomes with minimum percent identity cutoffs that may not sufficiently distinguish among sub-clades (83, 84). However, the apparent diversity within the Accumulibacter lineage may require reevaluation of the phylogeny and taxonomy overall. Although the *ppk1*-based clade designations coincide with genome-wide ANI based cutoffs remarkably well (Supplementary Figure 2B), there are some discrepancies. A few Accumulibacter genomes have been assembled that do not adhere to the *ppk1*-based population structure but fall within the Accumulibacter lineage based on a species tree of single-copy core genes (Supplementary Figure 2A), such as the proposed novel species ‘*Candidatus* Accumulibacter aalborgensis’ (85). Additionally, a few genomes preliminarily classified by the Genome Taxonomy Database (GTDB) as Accumulibacter branch outside of the established nomenclature and are quite divergent based on the species and *ppk1* phylogenies as well as genome-wide and *ppk1* sequence similarities (Figure 2B, Supplementary Figure 2). Overall, substantial work is needed to reevaluate the phylogenetic structure of the Accumulibacter lineage as a whole and to connect with signatures of homologous recombination and evolutionary trajectories within coexisting clades to understand the genetic boundaries and thus ecological interdependencies of individual clades or strains.

Although the boundaries of individual Accumulibacter clades or strains have not been entirely resolved to infer separate ecotypes, denitrification capabilities of specific clades have been repeatedly suggested to be a niche-differentiating feature (14, 23, 25). Previously, two separate bioreactors fed with either acetate or propionate as the sole carbon source were enriched with different Accumulibacter morphotypes, which were also linked to different denitrification capabilities (28). A study with a bioreactor highly enriched in clade IC inferred simultaneous use of oxygen, nitrite, and nitrate as electron acceptors under micro-aerobic conditions either due to metabolic flexibility or heterogeneity within IC members (23). The resident strain IC-UW-LDO also possessed the full suite of genes for denitrification and expressed them in microaerobic conditions (14). Coexisting Accumulibacter strains with differing denitrifying abilities have also been observed in a denitrifying EBPR bioreactor enriched in clade IA and clade IC (17, 27). The clade IA strain encoded genes for complete denitrification, whereas the co-occurring clade IC strain and other flanking community members did not (27). Additionally, Gao et al. performed an ancestral genome reconstruction analysis similar to Oyserman et al. 2017 (67) to specifically understand the flux of denitrification gene families (27). The *nir* gene cluster was identified as a core COG to all sampled Accumulibacter genomes, whereas the periplasmic nitrate reductase *napAGH* and nitrous oxide reductase *nosZDFL* clusters were core to only Type I Accumulibacter genomes, and inferred to have been lost for some Type II genomes (27). Additionally, Wang et al. subsequently observed that an internal stop codon within the open reading frame of the *nosZ* gene in a clade IC strain coincided with decreased transcripts mapping back (17, 27). In our study, we observed differential expression of denitrification genes in strain IA-UW4 but not strain IIC-UW6. These expression differences could explain why the two strains can coexist and contribute to overall ecosystem functioning (e.g. phosphorus removal) by using different electron acceptors in the anaerobic/aerobic conditions. Although the synthetic feed for the bioreactors in this study inhibits ammonia oxidation through allylthiourea addition (and thus inhibits nitrite/nitrate production to fuel respiratory denitrification), the expression profiles of denitrification genes in strain IA-UW4 suggests this strain is equipped to have a differentiating role in a denitrifying bioreactor. We specifically observed differential expression of nitrate reductase and nitrous-oxide reductase genes in strain IA-UW4, where homologs of these genes were not present in the strain IIC-UW6 genome and other Type II genomes. From our work and previous studies, lack of the complete genetic repertoire for denitrification among and within individual clades in addition to transcriptional-based evidence suggests that denitrification capabilities could be a niche differentiating role for specific Accumulibacter populations.

Accumulibacter’s anaerobic metabolism has also been shown to differ under varying substrate or stoichiometric conditions and possibly between the two Accumulibacter types. Recent simulations of ‘omics datasets paired with enzymatic assays hint that the routes of anaerobic metabolism for Accumulibacter may differ based on environmental conditions (73). Under high acetate conditions, polyphosphate accumulation arises through the glyoxylate shunt, whereas under increased glycogen concentrations, glycolysis is used in conjunction with the reductive branch of the TCA cycle (73). Furthermore, Welles et al. found that Type II Accumulibacter can switch to partial glycogen degradation more quickly than Type I, enabling Type II to fuel VFA uptake in polyphosphate-limited conditions (86, 87). Conversely, when concentrations of polyphosphate are high, VFA uptake rates are higher in Type I. However, these studies only identified Accumulibacter lineages based on *ppk1* types, and therefore extrapolated metabolic assumptions from presumably more resolved strains to the entire type. These broad interpretations may obscure apparent functional differences between the two types and individual clades. Future experiments could integrate multi-‘omics approaches to investigate not only metabolic flexibility under different substrate conditions and perturbations but also how metabolisms are distributed amongst Accumulibacter clades in bioreactor ecosystems that are not completely enriched in a single clade or strain and elucidate key functional differences.

## CONCLUSIONS

In this study, we integrated genome-resolved metagenomics and time-series metatranscriptomics to understand gene expression patterns of co-existing Accumulibacter strains within a bioreactor ecosystem. We found evidence for denitrification gene expression in the dominating IA-UW4 strain, but higher overall transcriptional activity for the IIC-UW6 strain along with differential expression of a high-affinity phosphorus transporter. The coexistence of Accumulibacter populations exhibiting different gene expression patterns for phosphorus and nitrogen cycling and hydrogen productions suggest niche differentiating features and contribution to different metabolic programs. Detailed experiments incorporating microautoradiography combined with fluorescence in situ hybridization (MAR-FISH) could be applied to confirm metabolic activities of coexisting Accumulibacter populations and how they contribute to important EBPR-related functions (88). These results also highlight the need to reevaluate the phylogenetic structure of the Accumulibacter lineage and define how potential genetic boundaries may translate to individual ecological niches that allow for the coexistence of multiple clades, species-like groups, or strains. Additionally, we highlight an approach for exploring functions of co-existing species-like uncultivated microbial lineages by exploring gene expression patterns of shared and accessory genes. Overall, this work emphasizes the need to understand the basic ecology and evolution of microorganisms underpinning biotechnological processes, which can lead to better treatment process control and resource recovery outcomes in the future (89).

## ACKNOWLEDGEMENTS

We thank Dr. Jeffrey Lewis for feedback on the differential expression analyses and interpretation. This work was supported by funding from the National Science Foundation (MCB-1518130) to K.D.M. and D.R.N. F.M. and P.C. received funding from the Chilean National Commission for Scientific and Technological Research (CONICYT). E.H.K. was supported by the 2017 UW-Madison Hilldale Scholarship. E.A.M. was supported by a University of Wisconsin – Madison Department of Bacteriology Fellowship. We thank the University of Wisconsin – Madison Biotechnology Center DNA Sequencing Facility for Illumina sequencing services. This research was performed in part using the Wisconsin Energy Institute computing cluster, which is supported by the Great Lakes Bioenergy Research Center as part of the U.S. Department of Energy Office of Science (DE-SC0018409).

---

**Supplementary Figure 1.**
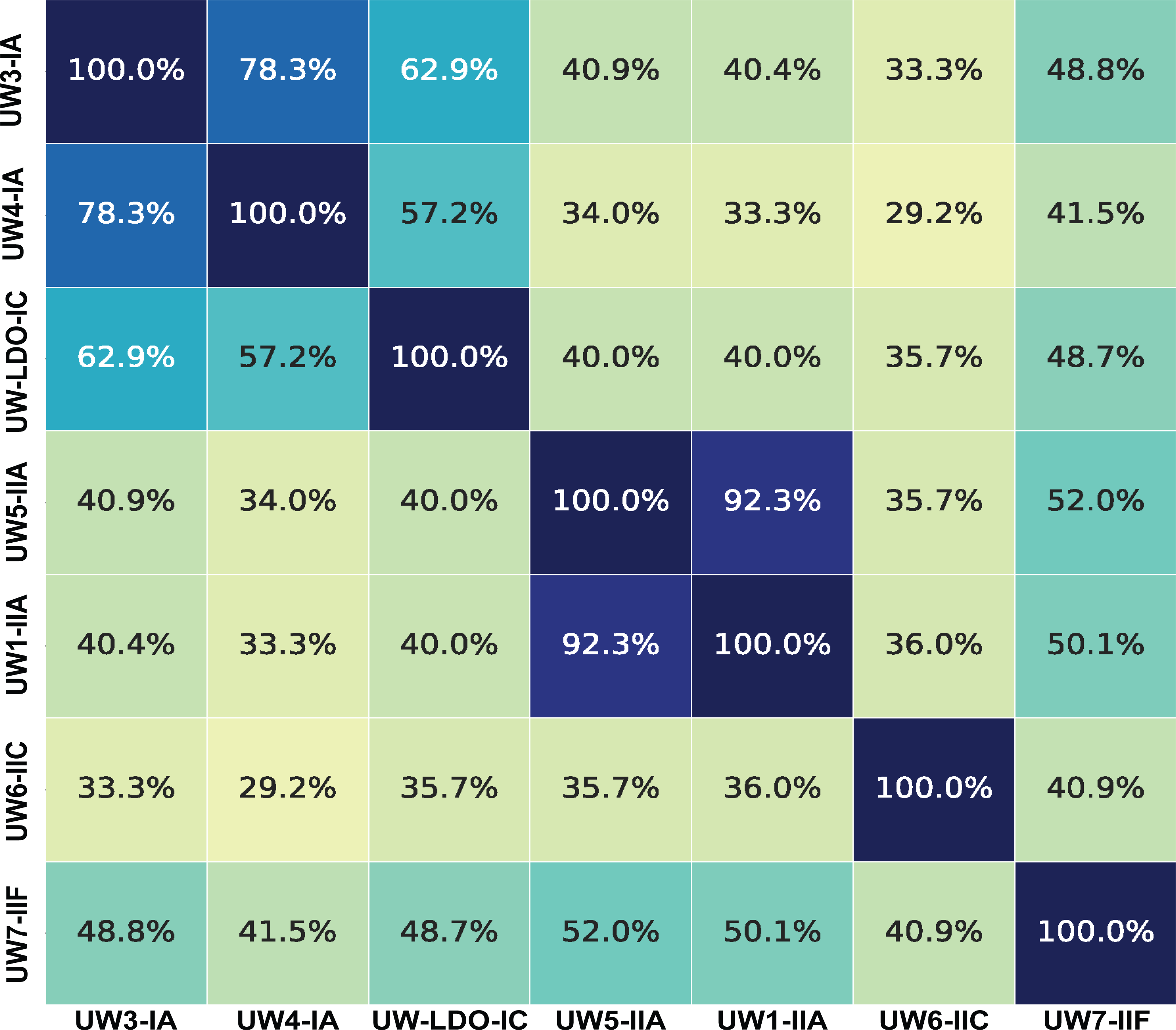
Pairwise Pearson correlation coefficient of shared gene content as a measure of how similar each of the UW genomes is to each other based on COGs.

**Supplementary Figure 2.**
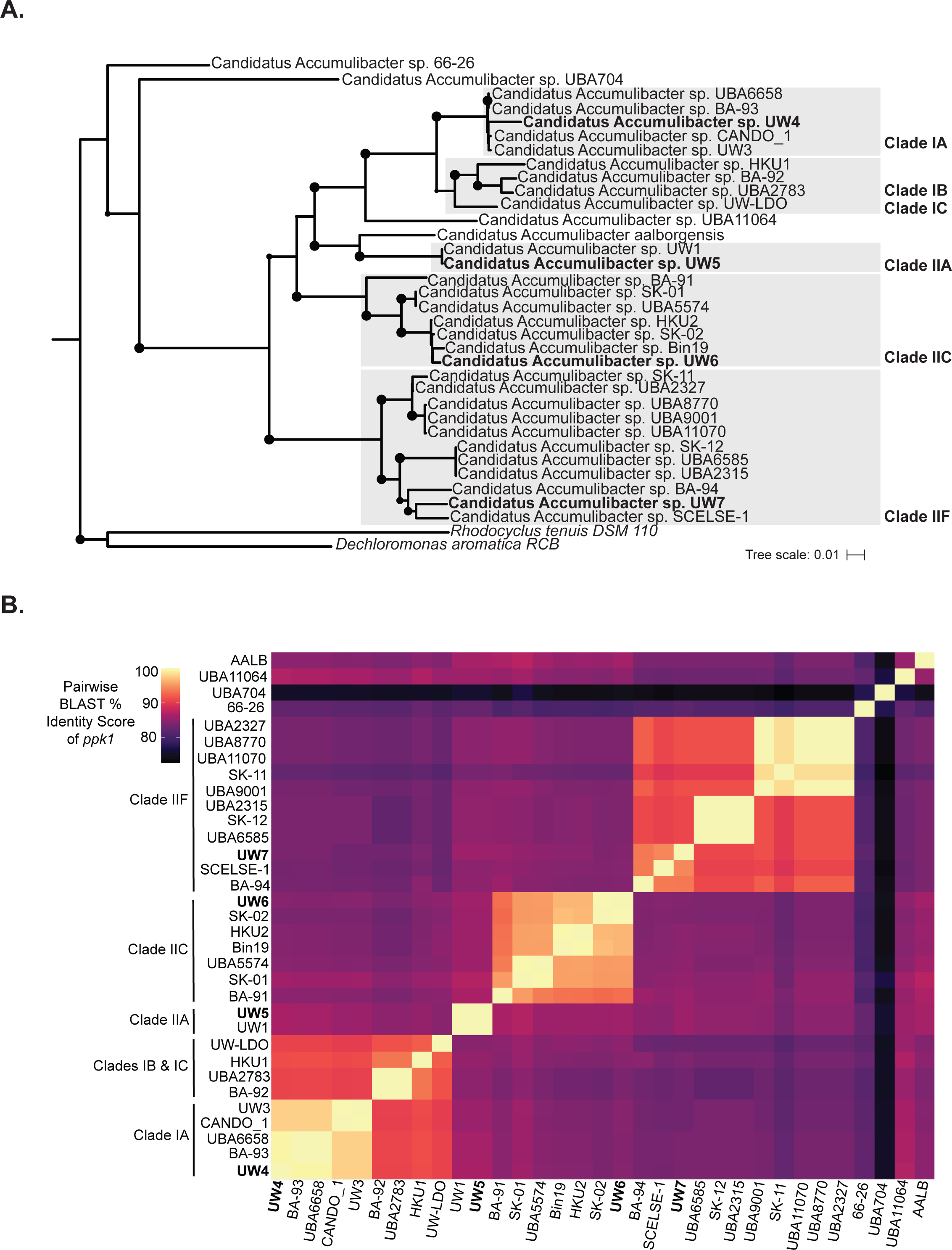
**S2A)** Species tree of all Accumulibacter reference genomes, UW-assembled genomes, and outgroup genomes using single-copy markers from the GTDB-tk (50). A phylogenetic tree was constructed using RAxmL with 100 rapid bootstraps and visualized in iTOL. **S2B)** Pairwise BLAST percent identity scores for *ppk1* coding regions for each reference genome in Figure 2B, in order of the *ppk1* phylogeny in Figure 2A. Raw pairwise percent identity scores are available at https://figshare.com/articles/dataset/Pairwise_Accumulibacter_ppk1_BLAST_Percent_D_Scores/13237247.

**Supplementary Table 1** Genome accessions and statistics for all Accumulibacter references used in phylogenies, ANI comparisons, and COG calculations. A reference genome was used for COG comparisons if it was a high quality genome (>95% complete and <5% redundant). Available at https://figshare.com/account/projects/90614/articles/13237148

**Supplementary File 1** Raw pairwise genome-wide ANI results for each genome calculated with FastANI. Available at https://figshare.com/account/projects/90614/articles/13237244

**Supplementary File 2** Raw pairwise BLAST percent identity (PID) results for *ppk1* genes from each reference genome. Available at https://figshare.com/account/projects/90614/articles/13237247

## Notes

https://figshare.com/projects/Metabolic_Plasticity_of_Accumulibacter_Clades/90614

### Competing Interest Statement

The authors have declared no competing interest.

### Summary of Updates

Minor modifications to manuscript text, figure fixes

